# Validation of a single-step, single-tube reverse transcription-loop-mediated isothermal amplification assay for rapid detection of SARS-CoV-2 RNA

**DOI:** 10.1101/2020.04.28.067363

**Authors:** Jean Y. H. Lee, Nickala Best, Julie McAuley, Jessica L. Porter, Torsten Seemann, Mark B. Schultz, Michelle Sait, Nicole Orlando, Karolina Mercoulia, Susan A. Ballard, Julian Druce, Thomas Tran, Mike G. Catton, Melinda J. Pryor, Huanhuan L. Cui, Angela Luttick, Sean McDonald, Arran Greenhalgh, Jason C. Kwong, Norelle L. Sherry, Maryza Graham, Tuyet Hoang, Marion Herisse, Sacha J. Pidot, Deborah A. Williamson, Benjamin P. Howden, Ian R. Monk, Timothy P. Stinear

## Abstract

2.

**Introduction:** The SARS-CoV-2 pandemic of 2020 has resulted in unparalleled requirements for RNA extraction kits and enzymes required for virus detection, leading to global shortages. This has necessitated the exploration of alternative diagnostic options to alleviate supply chain issues.

**Aim:** To establish and validate a reverse transcription loop-mediated isothermal amplification (RT-LAMP) assay for the detection of SARS-CoV-2 from nasopharyngeal swabs.

**Methodology:** We used a commercial RT-LAMP mastermix from OptiGene Ltd in combination with a primer set designed to detect the CDC N1 region of the SARS-CoV-2 nucleocapsid (N) gene. A single-tube, single-step fluorescence assay was implemented whereby as little as 1 μL of universal transport medium (UTM) directly from a nasopharyngeal swab could be used as template, bypassing the requirement for RNA purification. Amplification and detection could be conducted in any thermocycler capable of holding 65°C for 30 minutes and measure fluorescence in the FAM channel at one-minute intervals.

**Results:** Assay evaluation by assessment of 157 clinical specimens previously screened by E-gene RT-qPCR revealed assay sensitivity and specificity of 87% and 100%, respectively. Results were fast, with an average time-to-positive (Tp) for 93 clinical samples of 14 minutes (SD ±7 minutes). Using dilutions of SARS-CoV-2 virus spiked into UTM, we also evaluated assay performance against FDA guidelines for implementation of emergency-use diagnostics and established a limit-of-detection of 54 Tissue Culture Infectious Dose 50 per ml (TCID_50_ mL^−1^), with satisfactory assay sensitivity and specificity. A comparison of 20 clinical specimens between four laboratories showed excellent interlaboratory concordance; performing equally well on three different, commonly used thermocyclers, pointing to the robustness of the assay.

**Conclusion:** With a simplified workflow, N1-STOP-LAMP is a powerful, scalable option for specific and rapid detection of SARS-CoV-2 and an additional resource in the diagnostic armamentarium against COVID-19.

**Data summary:** The authors confirm all supporting data, code and protocols have been provided within the article or through supplementary data files.

## 4. Introduction

The current SARS-CoV-2 pandemic has created an unprecedented global demand for rapid diagnostic testing. Most countries are employing real-time quantitative PCR (RT-qPCR) for confirmation of infection [1-7]. Consequently, a global shortage of RNA extraction kits as well as RT-qPCR assay kits and their associated reagents has ensued [8-10]. Therefore, alternative diagnostics not dependent on these commonly used materials are required. First described 20 years ago by Notomi *et al*., the loop-mediated isothermal amplification (LAMP) assay is robust, rapid and straightforward, yet retains high sensitivity and specificity [11]. These features have seen the LAMP assay and the inclusion of a reverse transcriptase (RT-LAMP) implemented for a broad range of molecular diagnostic applications extending from infectious diseases, including detection of the original SARS-CoV-1 virus [12], bacteria and parasites [13-15] to cancer [16]. The advantages of RT-LAMP include; using different reagents than RT-qPCR, the potential for direct processing of samples without the need for prior RNA extraction and an extremely rapid turn-around time. Several groups have now described different RT-LAMP assays for detection of SARS-CoV-2 RNA [17-24].

In this study, we developed a RT-LAMP assay that targets the CDC N1 region of the SARS-CoV-2 nucleocapsid gene (N-gene) [3] and used a commercial mastermix from OptiGene Ltd. This mix contains a proprietary reverse transcriptase for cDNA synthesis and the thermophilic GspSSD strand-displacing polymerase/reverse transcriptase for DNA amplification (www.optigene.co.uk) with a dsDNA intercalating fluorescent dye. Detection was achieved by measuring the increase in fluorescence as amplification products accumulate. The **N1** gene **S**ingle **T**ube **Op**tigene **LAMP** assay (hereafter called N1-STOP-LAMP) was assessed for the direct detection of SARS-CoV-2 RNA, following *FDA Policy for Diagnostic Tests for Coronavirus Disease-2019 during the Public Health Emergency* against the four parameters and acceptance criteria summarised in Table 1 [5]. Validation samples were human upper respiratory tract specimens, collected using nasopharyngeal flocked swabs stored in universal transport media (UTM).

**Table 1.** FDA validation study recommendations for SARS-CoV-2 molecular diagnostics [5].

| Method Validation Parameter | Validation description and acceptance criteria |
| --- | --- |
| 1. Limit of Detection (LoD) | Test a dilution series of three replicates per concentration. LoD defined as lowest concentration at which 19/20 replicates positive. |
| 2. Clinical evaluation | Test at least 30 contrived reactive specimens and 30 non-reactive specimens. At least 20 specimens should have virus/RNA concentration of 1-2x LoD, remainder spanning the assay range. Acceptance defined as 95% agreement 1-2x LoD, and 100% agreement at other concentrations and for negative specimens. |
| 3. Inclusivity | *In silico* analysis indicating percent identity matches of target region against publicly available SARS-CoV-2 sequences. Acceptance defined as 100% match against all entries with the selected primers/probes. |
| 4. Cross-reactivity | *In silico* analysis of the assay primers/probes compared to respiratory microbiota and other viral pathogens. Acceptance defined as <80% homology between all primers/probes and non-target sequences present in public databases. |

## 5. Methods

### Specimen collection and handling

Nasopharyngeal swabs were collected by qualified healthcare professionals from patients meeting the epidemiological and clinical criteria as specified by the Victorian Department of Health and Human Services at the time of swab collection [1] between the 23^rd^ of March 2020 and 4^th^ of April 2020. Copan flocked swabs collected and directly inoculated on site in either 1 mL or 3 mL of UTM® (Catalogue N°s., 330C and 350C respectively) were used. Samples were collected at metropolitan hospitals in Melbourne, Victoria, Australia, and transported to the Doherty Institute Public Health Laboratories for further testing as per World Health Organization recommendations [6]. All swabs were processed in a class II biological safety cabinet.

### Cell culture and SARS-CoV-2 RNA extraction

Vero cells (within 30 passages from the original American Type Culture Collection [ATCC] stock) were maintained in Minimal Essential Media (MEM) supplemented with 10% heat-inactivated foetal bovine serum (FBS), 10 μM HEPES, 2 mM glutamine and antibiotics. Cell cultures were maintained at 37°C in a 5% CO_2_ incubator. All virus infection cultures were conducted within the High Containment Facilities in a PC3 laboratory at the Doherty Institute. To generate stocks of SARS-CoV-2, confluent Vero cell monolayers were washed once with MEM without FBS (infection media) then infected with a known amount SARS-CoV-2 virus originally isolated from a patient [25]. After 1h incubation at 37°C in a 5% CO2 incubator to enable virus binding, infection media containing 1 mg mL^−1^ TPCK-trypsin was added, and flasks returned to the incubator. After 3d incubation and microscopic confirmation of widespread cytopathic effect (CPE), the supernatants were harvested, and filter sterilised through a 0.45 μm syringe filter. To assess infectious SARS-CoV-2 viral titres, both Tissue Culture Infectious Dose 50 (TCID_50_) and plaque assays were performed. Briefly, serial dilutions of the stock virus were added to washed monolayers of Vero cells. After 1h incubation to allow virus to adhere, for the TCID_50_ assay, infection media containing 1 μg mL^−1^ TPCK-Trypsin was added, while for the plaque assay the infected cell monolayer was overlaid with Leibovitz-15 (L15) media supplemented with 0.9% agarose (DNA grade, Sigma), antibiotics and 2 μg mL^−1^ TPCK Trypsin. After 3d incubation, the dilution of stock required to cause CPE in at least 50% of wells (TCID_50_) was determined via back calculation of microscopic confirmation of CPE in wells for a given dilution. For quantitation via plaque assay, plaques present in the monolayer were macroscopically visualised, individually counted and back calculated for each given dilution to determine the number of Plaque Forming Units (PFU) per mL in the original stock. Stocks of SARS-CoV-2 used by this study had a TCID_50_ of 5.4×10^5^ mL^−1^ and plaque assay gave 9.78×10^5^ PFU mL^−1^. To heat inactivate the virus, 200 μL of neat stock was heated to 60°C for 30 min, then cooled. Inactivation was confirmed via complete lack of CPE and plaque formation using both TCID_50_ and plaque assays. To prepare SARS-CoV-2 RNA from stocks, 500 μL aliquots were thawed and RNA extracted using the RNeasy mini Kit (Qiagen) according to the manufacturer’s specifications, with an elution volume of 50 μL. Based on RNA concentrations, the total virus harvested from Vero cell culture was 3.1×10^9^ copies mL^−1^, suggesting a high number of non-infectious virus particles in the virus stocks.

### RNA extraction from UTM for RT-qPCR

A 200 μL aliquot of Copan UTM® was processed through the QIAsymphony DSP Virus/Pathogen Mini Kit (Qiagen, Cat# 937036) following manufacturer’s instructions on the QIAsymphony SP instrument. The RNA was eluted in 60 μL of recommended buffer.

### RNA extraction from UTM SPRI beads for N1-STOP-LAMP

Solid-phase reversible immobilisation (SPRI) on carboxylated paramagnetic beads (Sera-Mag™ Magnetic SpeedBeads™, from GE Healthcare) were prepared for RNA binding as described (https://openwetware.org/wiki/SPRI_bead_mix). RNA purification was performed in 96-well plates with initial lysis by the addition of 25 μL of 6GTD lysis buffer (7.08 g 10 mL^−1^ guanidine thiocyanate (6M), made up to 8.2 mL with water, 1 mL Tris HCL [pH 8.0] and 800 μL of 1M dithiothreitol), mixed by pipetting ten times and incubation at room temperature for 1 min [26]. To this, 75 μL of 100% ethyl alcohol and 20 μL of prepared SPRI beads were added, mixed by pipetting ten times and incubated at room temperature for 5 min. The RNA-bead complex was then immobilised by placing the 96-well plate on a magnetic rack and incubated again at room temperature for 5 min. The supernatant was then discarded, and the beads washed twice in 200 μL of freshly prepared 80% ethyl alcohol (v/v) with 30 sec room temperature incubation between each wash. The beads were air-dried for 2 min at room temperature before RNA was eluted by the addition of 20 μL of nuclease-free water.

### E-gene RT-qPCR

A one-step RT-qPCR was conducted with the primers as described by Corman *et al*., targeting the viral envelope E-gene of SARS-CoV-2 E_Sarbeco_F: 5’-ACAGGTACGTTAATAGTTAATAGCGT-3’, 5’-E_Sarbeco_R: ATATTGCAGCAGTACGCACACA-3’, E_Sarbeco_P: 5’-FAM-ACACTAGCCATCCTTACTGCGCTTCG-BHQ1-3’) [27]. A 20 μL reaction was assembled consisting of 1x qScript XLT One-Step RT-qPCR ToughMix Low ROX (2x) (QuantaBio), 400 nM E_Sarbeco_F, 400 nM E_Sarbeco_R 200 nM E_Sarbeco_P (Probe) and 5 μL of purified RNA. The following program was conducted on an ABI 7500 Fast instrument: 55°C for 10 min for reverse transcription, 1 cycle of 95°C for 3 min and then 45 cycles of 95°C for 15 sec and 58°C for 30 sec.

### N1-STOP-LAMP

A 50-reaction bottle of dried Reverse Transcriptase Isothermal Mastermix (Optigene, ISO-DR004-RT) was rehydrated with 750 μL of resuspension buffer and vortexed gently to mix. The mix contains both a proprietary reverse transcriptase and the GspSSD LF DNA polymerase that has both reverse transcriptase and strand-displacing DNA polymerase activity. The N1-STOP-LAMP assay uses six standard LAMP oligonucleotide primers that target the sequence spanning the CDC N1 region of the SARS-CoV-2 nucleocapsid gene. Sequences of the primers are available upon request (GeneWorks Ltd). Each N1-STOP-LAMP reaction contained: 15 μL of mastermix, 5 μL of 5x primer stock and 4 μL of water. Mastermix was reconstituted in a separate biological safety cabinet to that used for template addition. The source of RNA template for the reaction consisted of either 1 μL of purified SARS-CoV-2 RNA, 1 μL of SARS-CoV-2 purified virus or 1 μL of UTM from a nasopharyngeal swab. A no-template control (1 μL water) was included in all runs. Reactions were assembled in either 8-tube Genie strips (OP-00008, OptiGene Ltd) or 96-MicroAmp-Fast-Optical reaction plate (Applied Biosystems). Strip tubes reactions were capped, or for 96-MicroAmp-Fast-Optical reaction plates, sealed with MicroAmp Optical adhesive film (Applied Biosystems) and tapped to remove bubbles. Strip tubes were loaded onto the Genie-II or Genie-III (OptiGene Ltd) and 96-well plates run on a QuantStudio 7 thermocycler (ThermoFisher). Reactions were incubated at 65°C for 30 min with fluorescence acquisition every 30 sec (Genie instruments) or 1 min (QuantStudio 7). A positive result was indicated by an increase in fluorescence at an emission wavelength of 540 nm (FAM channel) above a defined threshold, recorded as time-to-positive (Tp) expressed in min:sec.

### Dilution series of purified SARS-CoV-2

A virus stock with of 5.4 x10^5^ TCID_50_ mL^−1^ was serially diluted to 10^−6^ in a generic UTM (comprising per liter: 8 g NaCl, 0.2 g KCl, 1.15 g Na_2_HPO_4_, 0.2 g KH_2_PO_4_, 4 mL 0.5% phenol red 0.5%, 5 g gelatin, 950 mL tissue culture water, 1 mL Fungizone [5 mg mL^−1^], 20 mL penicillin/streptomycin) in biological triplicates, with the 10^−1^ through to the 10^−6^ dilutions tested by N1-STOP-LAMP (1 μL). RNA was extracted from a 200 μL aliquot of each dilution as described above and 5 μL of the purified RNA used as template in E-gene RT-qPCR or N1-STOP-LAMP assay.

### Interlaboratory comparison of N1-STOP-LAMP

A panel of 20 blinded clinical samples (13 positive and 7 negative) with cycle threshold (Ct) values previously established by E-gene RT-qPCR were aliquoted and distributed to three different laboratories for independent testing by N1-STOP-LAMP assay (Table S2).

### Biological specificity

To test for cross reactivity of the N1-STOP-LAMP assay, a control panel of respiratory pathogens (NATRPC2-BIO, ZeptoMetrix) was screened (Table S3). A 1 μL aliquot of NATrol RP1 or RP2, or samples spiked with SARS-CoV-2 virus were used as template for N1-STOP-LAMP.

### In silico nucleotide sequence comparisons of the N1 region of SARS-CoV-2

To assess the inclusivity and exclusivity of the N1 region targeted by the LAMP assay 2755 publicly available SARS-CoV-2 genomes were downloaded and filtered to remove entries of less than 29 kb using *Seqtk seq* (v1.3-r106). Sequence gaps were replaced with ‘N’ using *Seqkit* (v0.12.0). After this quality control step, homology searches were then conducted using *NCBI BLAST+ blastn* using *Genepuller*.*pl* (https://github.com/tseemann/bioinfo-scripts/blob/master/bin/gene-puller.pl) to find the region in each of the remaining 2738 genomes matching the 5’ region of the N1 sequence. The resulting sequence coordinates of those hits were then used to extract the 240 bp region from all 2738 genomes using *Genepuller*.*pl* and aligned with Clustalo (v1.2.4). The alignment was visualised with *Mesquite* (v3.61).

### Statistical analysis

Data analysis was managed using GraphPad Prism (v8.4.1). Sensitivity and specificity testing were performed using the Wilson/Brown hybrid method as deployed in GraphPad Prism.

## 6. Results

### Assessing detection sensitivity of N1-STOP-LAMP using purified RNA

We began by assessing the limit of detection (LoD) of N1-STOP-LAMP under optimum conditions using a 10-fold dilution series of purified RNA, prepared from a titred SARS-CoV-2 virus stock. Although LAMP assays are not strictly quantitative, a positive correlation between Tp and RNA concentration was evident. This was indicated by an absolute detection threshold between 50 and 500 viral genome copies per reaction. We achieved a reliable detection of 5/5 replicates at 500 viral genome copies and detection of 4/5 replicates at 50 viral genome copies per reaction (Fig. 1A). Of note, this threshold is likely an underestimate of the true detection limit, as we assumed all RNA yielded from the viral stock generated from the Vero cell supernatant was of viral origin. Using TCID_50_, the absolute LoD of N1-STOP-LAMP was between 0.001 and 0.01 TCID_50_ per reaction (equivalent to 1-10 TCID_50_ mL^−1^).

**Fig. 1.**
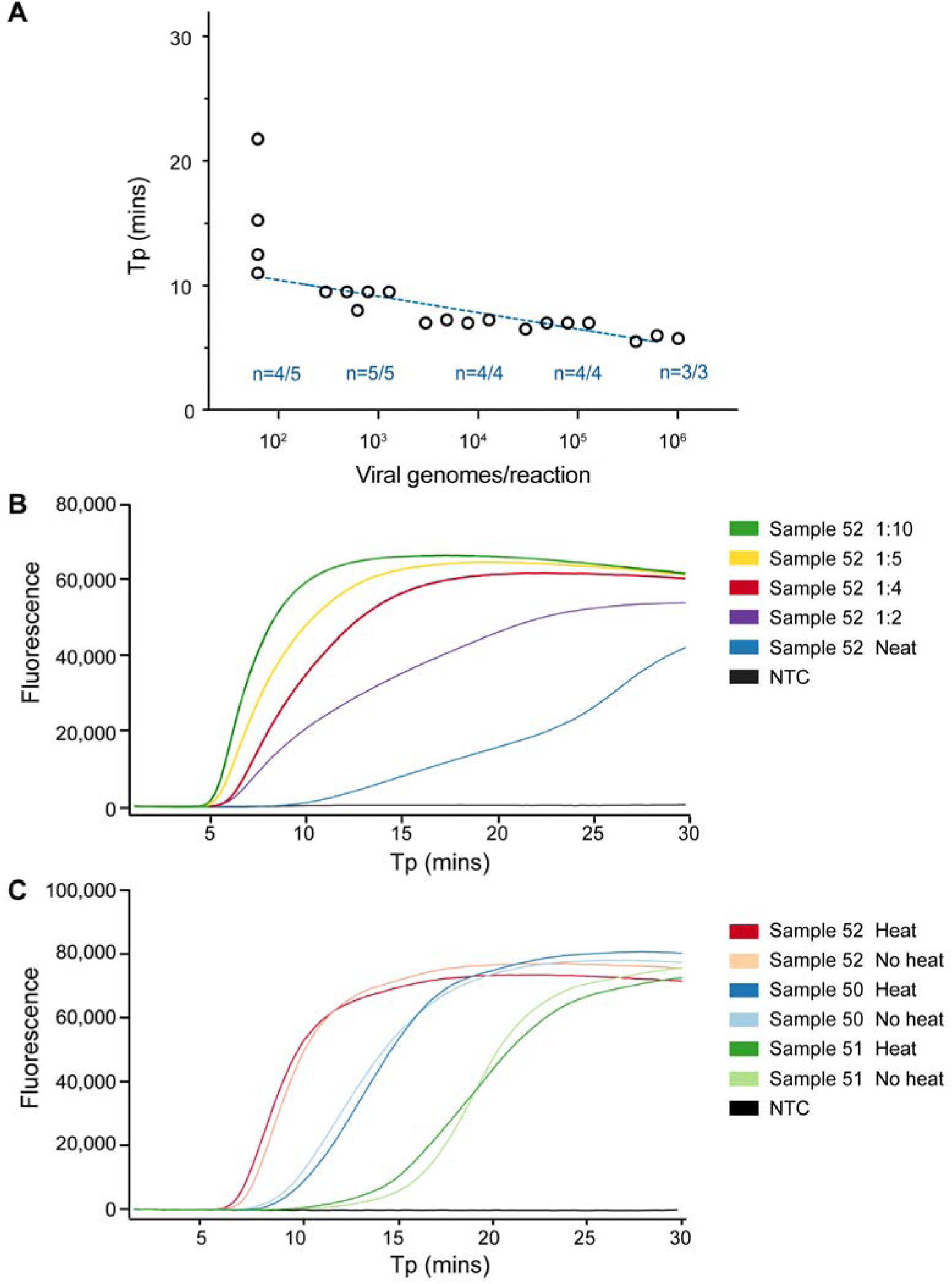
Limit of detection of N1 STOP-LAMP and establishing optimal sample template volume and inactivation conditions. (A) Plot showing performance of the N1 LAMP assay across 5-log_10_ dilution of purified viral RNA. Y-axis is time-to-positive (Tp) and x-axis is estimate of viral genomes/reaction based on starting RNA concentration. The number of replicates per dilution (n) and number of positive replicates per dilution is indicated. (B) Example N1-STOP-LAMP amplification plots for SARS-CoV-2 positive clinical sample N°. 52, showing inhibitory impact of 5 μL of a neat sample matrix on LAMP and the effect of diluting the UTM in water. (C) N1-STOP-LAMP amplification plots for three SARS-CoV-2 positive clinical samples, showing no loss in detection sensitivity after specimen heat-treatment of 60°C for 30 min to inactivate virus.

### Direct detection of SARS-CoV-2 RNA in clinical specimens and specimen inactivation

The nasopharyngeal specimens received by our public health laboratories were collected using Copan flocked swabs in either 1 mL or 3 mL of UTM®. To assess the impact of sample matrix on the LAMP assay, we performed pilot experiments with UTM from swabs taken from SARS-CoV-2 negative specimens, adding increasing amounts of UTM to N1-STOP-LAMP. Sterile UTM had no impact on N1-STOP-LAMP (data not shown). However, the nasopharyngeal secretions in patient samples were observed to be inhibitory. Most pronounced when 5 μL of patient sample was used as the direct RNA template (Fig. 1B), but relieved at a 1/5 dilution of the patient sample, thus 5 μL of a 1/5 dilution of the patient sample or a 1 μL of neat patient sample was selected as the optimum template volume for N1-STOP-LAMP (Fig. 2A). Experiments were also conducted using dry swabs eluted in PBS. We tested elution in 0.5 mL, 1.0 mL and 1.5 mL and found that an elution volume of 1.5 mL of PBS was a good compromise between unnecessary dilution of potential virus in the sample and sufficient dilution to alleviate assay inhibition (data not shown). To reduce the risk associated with handling clinical specimens containing infectious SARS-CoV-2, we also assessed the impact of a heat inactivation step on detection sensitivity. Pre-treatment at 60°C for 30 min led to a >5-log_10_ inactivation of the virus accessed by PFU and TCID_50_ determination (data not shown). No impact on N1-STOP-LAMP detection sensitivity was observed, with equivalent Tp between each treatment (Table 2, Fig. 1C).

**Table 2.**
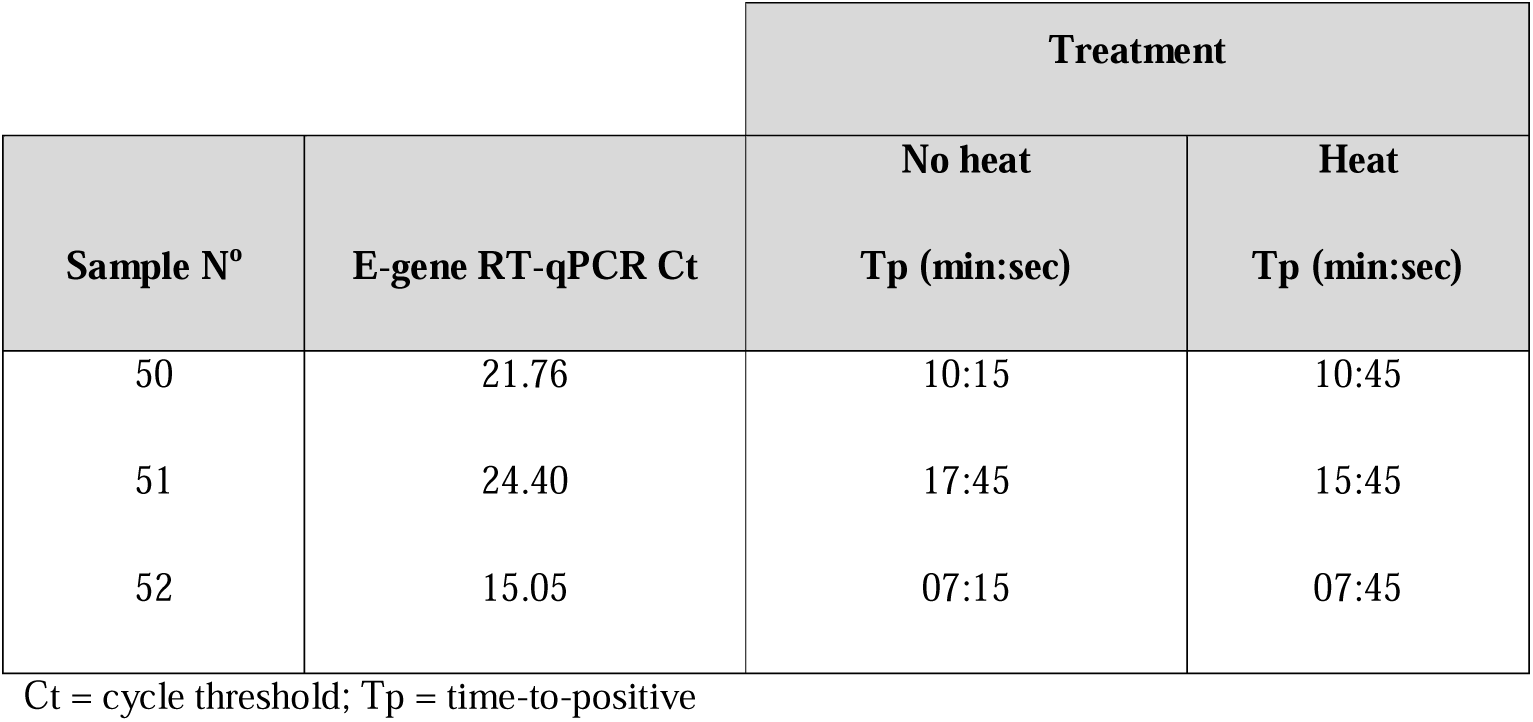
Assessment of heat treatment of universal transport medium specimens on detection sensitivity.

**Fig. 2:**
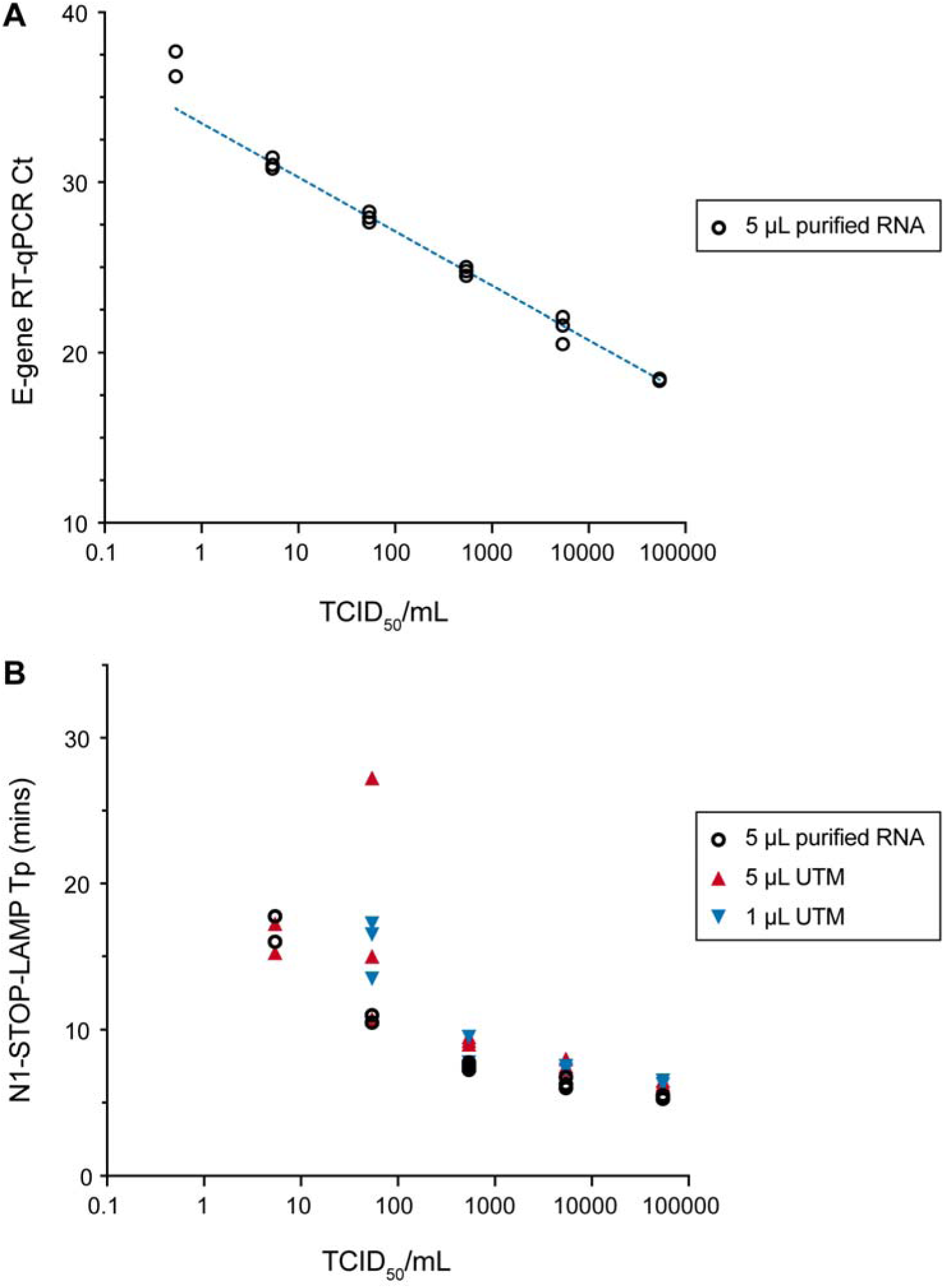
Comparison of E-gene RT-qPCR with N1-STOP-LAMP and limit of detection. Titred virus was serially diluted 10-fold in UTM in triplicate and RNA was extracted from each replicate dilution. A 1 μL aliquot of each UTM dilution was also tested directly by N1-STOP-LAMP. (A) Shows calibration curve for the E-gene RT-qPCR. A curve was interpolated using linear regression. (B) Comparative performance of N1-STOP-LAMP with 5 μL purified RNA versus 1 μL or 5 μL of UTM directly added to the reaction.

### Comparative detection sensitivity of N1-STOP-LAMP versus E-gene RT-qPCR

The N1-STOP-LAMP assay was then directly compared with E-gene RT-qPCR, using a dilution of titred SARS-CoV-2 virus stock in UTM across a 6-log_10_ dilution series. A RT-qPCR calibration curve was established from triplicate extraction experiments for each of the six dilutions (Fig. 2A). The assays were prepared with 5 μL of purified viral RNA as template for both the N1-STOP-LAMP and RT-qPCR assays. A 1 μL and 5 μL aliquot of each virus dilution in UTM was also tested directly in the N1-STOP-LAMP assay (*i*.*e*. without RNA extraction). The side-by-side comparisons showed E-gene RT-qPCR was up to 1-log_10_ more sensitive than N1-STOP-LAMP, with RT-qPCR detecting 2/3 replicates at the lowest dilution of 0.54 TCID_50_ mL^−1^ and N1-STOP-LAMP detecting no viral RNA at this concentration (Table 3, Fig. 2). At 54 TCID_50_ mL^−1^, the RT-qPCR and N1-STOP-LAMP detected 3/3 replicates. There was no difference in N1-STOP-LAMP sensitivity using either 5 μL UTM added directly to the test or purified RNA, but the assay lost another 1-log_10_ sensitivity where 1 μL of neat UTM was added directly to the N1-STOP-LAMP reaction (Fig. 2B).

**Table 3.** Comparison of N1-STOP-LAMP and RT-qPCR performance using titred virus diluted in universal transport medium.

| SARS-CoV-2<br>(TCID<sub>50</sub> mL<sup>-1</sup>) | N | N1-LAMP<br>(1 µL direct) |  | N1-LAMP<br>(5 µL direct) |  | N1-LAMP<br>(5 µL of 60 µL) |  | RT-qPCR<br>(5 µL of 60 µL) |  |
| --- | --- | --- | --- | --- | --- | --- | --- | --- | --- |
|  |  | Tp<br>(Avg) | CV<br>(%) | Tp<br>(Avg) | CV<br>(%) | Tp<br>(Avg) | CV<br>(%) | Ct<br>(Avg) | CV<br>(%) |
| 54000 | 3 | 06.33 | 02.28 | 06.41 | 02.25 | 05.33 | 02.71 | 18.41 | 0.33 |
| 5400 | 3 | 07.42 | 01.95 | 07.58 | 05.04 | 06.33 | 06.03 | 21.40 | 3.75 |
| 540 | 3 | 08.92 | 11.33 | 09.25 | 02.70 | 07.50 | 03.33 | 24.77 | 1.03 |
| 54 | 3 | 15.75 | 12.60 | 17.67 | 48.49 | 10.67 | 02.71 | 27.94 | 1.13 |
| 5.4 | 3 | nd | - | 16.25 | 08.70<sup>#</sup> | 16.88 | 07.33<sup>#</sup> | 31.10 | 1.17 |
| 0.54 | 3 | nd | - | nd | nd | nd | - | 36.96 | 2.79<sup>#</sup> |

Tp = time-to-positive; CV = Coefficient of variation; nd = not detected; <sup>#</sup> 2/3 replicates positive at this dilution.

### Establishing N1-STOP-LAMP limit of detection (LoD)

The FDA guidelines for implementation of emergency-use diagnostics defines LoD as the lowest concentration at which 19/20 replicates are positive (Table 1). Informed by the previous experiment (Fig. 2B), we selected a SARS-CoV-2 concentration of 54 TCID_50_ mL^−1^ (in UTM) and tested 20x 1 μL aliquots by N1-STOP-LAMP. We observed 20/20 positive reactions with an average Tp of 15.8 mins, inter-quartile range 12-19 mins and coefficient of variation of 31.18% (Fig. 3A).

**Fig. 3:**
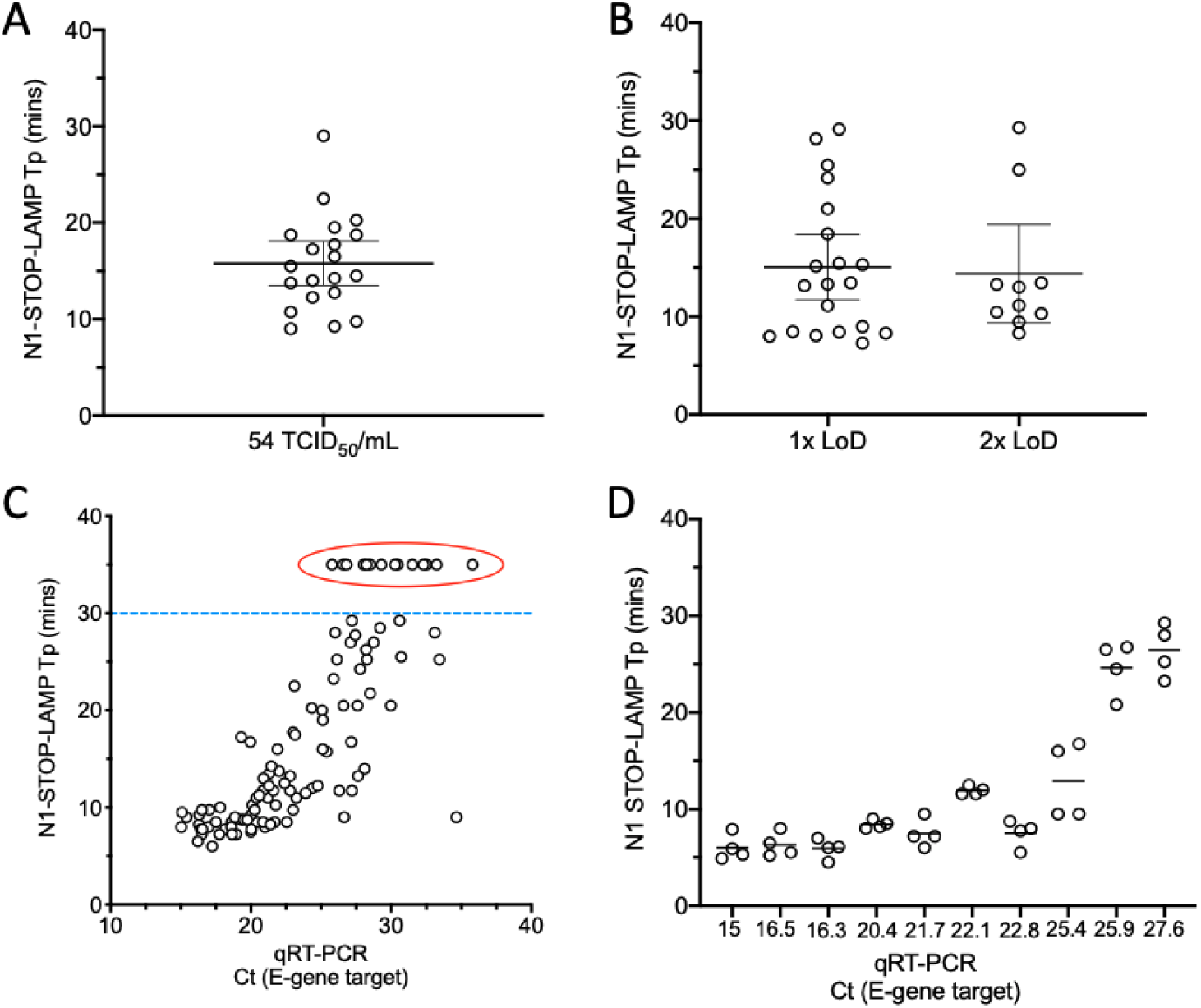
Clinical evaluation of N1-STOP-LAMP. (A) Establishing limit of detection according to FDA Emergency Use Authorisations (EUA) guidelines. Twenty, 1 μL replicates of SARS-CoV-2 virus in Universal Transport Medium (UTM) at a concentration of 54 TCID_50_ mL^−1^ were tested by direct N1-STOP-LAMP. All data points plotted, with average and 95% CI indicated. (B) Assessment of assay against FDA EUA clinical performance criteria. Thirty healthy volunteer dry nasopharyngeal swabs eluted in 1.5 mL of PBS were screened by N1-STOP-LAMP. Shown are time-to-positives (Tp). Mean and 95% CI are indicated. (C). Plot showing correspondence between 107 E-gene RT-qPCR positive and N1-STOP-LAMP, where 1 μL of clinical specimen in UTM was added directly to the RT-LAMP reaction. Y-axis is LAMP Tp and x-axis is RT-qPCR cycle threshold (Ct). The dotted line indicates Tp 30 mins. Results plotted above this line (encircled) were RT-qPCR positive but N1-STOP-LAMP negative. (D) Results of 4-way interlaboratory comparison of N1-STOP-LAMP. Ten clinical samples previously tested positive by E-gene RT-qPCR were used to create a test panel for comparing assay performance between laboratories.

### Evaluation of N1-STOP-LAMP against FDA criteria

The FDA guidelines for implementation of emergency-use diagnostics also require an assessment of assay performance using at least 30 contrived positive and 30 negative clinical specimens (authentic or contrived), with at least 20 of the positives at a concentration of 1-2x the LoD (Table 1). We obtained 30 nasopharyngeal dry swabs from healthy, anonymous volunteers. Swabs were eluted in 1.5 mL of PBS and 1 μL aliquots were tested directly by N1-STOP-LAMP. All eluates were negative. We then spiked the eluates with SARS-CoV-2 virus at 1x and 2x the LoD (approx. 50 and 100 TCID_50_ mL^−1^, respectively). Results showed that N1-STOP-LAMP detected 20/20 samples spiked with 1x the assay LoD and 10/10 samples at 2x LoD (Fig. 3B, Table 4). N1-STOP-LAMP thus meets the FDA Clinical Evaluation criteria (Table 1).

**Table 4.** Summary of clinical evaluation comparisons against FDA criteria.

|  | Negative | 1-2x LoD | 2x LoD |
| --- | --- | --- | --- |
|  |  | Tp (min:sec) | Tp (min:sec) |
| **N1-STOP-LAMP +** | 0 | 20 (07:30 – 22:00)<sup>#</sup> | 10 (08:30 – 21:00)<sup>#</sup> |
| **N1-STOP-LAMP -** | 30 | 0 | 0 |

LoD = Limit of detection; Tp = time-to-positive; <sup>#</sup> range indicated within parentheses.

### Clinical specimen evaluation of N1-STOP-LAMP against E-gene RT-qPCR gold standard

We then sought to evaluate N1-STOP-LAMP performance against E-gene RT-qPCR, setting the latter assay as the current ‘gold standard’. We directly screened 50 negative clinical specimens and 107 positive nasopharyngeal clinical specimens as established by E-gene RT-qPCR (Table S1). Results showed N1-STOP-LAMP assay sensitivity of 87%, specificity of 100%, positive predictive value of 100% and negative predictive value of 78% (Table 5, Fig. 3). As noted during initial sensitivity experiments (Table 3), a positive correlation was observed between N1-STOP-LAMP Tp and E-gene RT-qPCR Ct (Fig. 4). The average Tp among the 93 N1-LAMP-STOP positive specimens was 14 min (SD ± 7 min). For the false-negative samples with sufficient clinical material remaining (5 out of 14), RNA extraction using a simple, rapid magnetic bead purification protocol (takes 20 min) was performed, then N1-STOP-LAMP repeated. Four of the 5 samples returned a positive result with Tp range from 10:00 – 25:30 min:sec (Table S1). Indicating that the reduced sensitivity of N1-STOP-LAMP compared to RT-qPCR can be improved upon by using samples with a higher RNA template concentration and purity with only a small increase in time to results.

**Table 5.** Comparison of N1-STOP-LAMP versus E-gene RT-qPCR using clinical specimens.

|  | RT-qPCR + | RT-qPCR - | Total |
| --- | --- | --- | --- |
| N1-STOP-LAMP + | 93 | 0 | 93 |
| N1-STOP-LAMP - | 14 | 50 | 64 |
| **Total** | 107 | 50 | 157 |

### Confirmation of N1-STOP-LAMP inclusivity and exclusivity

The FDA guidelines for implementation of emergency-use diagnostics require an assessment of inclusivity and exclusivity of the oligonucleotide primers used in an assay. To assess the former, we performed *in silico* screening of the region spanned by the six primers used for the N1-STOP-LAMP assay against all available SARS-CoV-2 genome sequences (Supplementary data file 1). This 240 bp sequence spans the CDC N1 region of the SARS-CoV-2 N-gene [3]. Alignment of this region against 2738 genomes revealed 100% conservation amongst all entries, showing that this a conserved target sequence and that the primers used will perform equally across all identified lineages of the virus.

To assess the exclusivity criterion, we used a nucleotide BLAST search of the N1 region against the NCBI Genbank *nt* database and observed no non-SARS-CoV-2 sequence matches above 80% nucleotide identity, in-line with FDA cross-reactivity requirement. Further to this analysis, we also performed *in vitro* testing with the NATRPC2-BIO (ZeptoMetrix) specificity panel, a control panel consisting of 22 common respiratory pathogens. None of these pathogens were detected by N1-STOP-LAMP (Table S3). Thus, both *in silico* and *in vitro* testing confirmed the specificity of N1-STOP-LAMP for SARS-CoV-2.

### Interlaboratory comparisons

In order to explore the ability of other laboratories to readily deploy N1-STOP-LAMP, we prepared an external-quality-assessment (EQA) panel of 20 clinical specimens (Table S2). These were previous positive and negative clinical samples as assessed by E-gene RT-qPCR that had been heat-inactivated for safety. These specimens covered a range of virus titres (Ct values 16.3 – 30.6). The panel was tested simultaneously in four different laboratories (Labs A – D). The results showed excellent correspondence between all laboratories for ten positive samples, the majority with E-gene RT-qPCR Ct values <26 (Fig. 5, Table S2). Above this Ct (*i*.*e*. those samples with lower virus titres), the laboratories returned variably positive results (Samples 14, 16 & 18), reflecting that these specimens had virus concentrations at or beyond the N1-STOP-LAMP LoD (Table S2). Concordance among the seven negative results was overall very good, however one laboratory (Lab C) returned a false positive result (Sample 2), highlighting the issue of contamination with sensitive molecular tests (Table S2). Of note, the laboratories used a range of different instruments for amplification/detection including: OptiGene Genie II and III, BioRad CFX and ThermoFisher Quantstudio 7. This trial suggests that N1-STOP-LAMP is a robust and transferrable assay format.

## 7. Discussion

A critical component of an effective COVID-19 pandemic response is the rapid and robust detection of positive clinical samples [28]. This has required massively upscaled SARS-CoV-2 diagnostic capabilities. Particularly, a worldwide focus on developing improved technologies [29]. There have been more than 20 molecular tests recently receiving FDA Emergency Use Authorisations (EUAs) [5]. In this current study, we demonstrate the potential of N1-STOP-LAMP. We focused efforts on the LAMP assay because it has been previously employed for the rapid and robust detection of numerous RNA viruses including Zika, Chikungunya, Influenza and SARS-CoV-1 [12, 30-33] and was of great assistance during these outbreaks, particularly in resource constrained settings. Advantages of LAMP testing include rapid turnaround time, ease of implementation, non-standard reagent use and potential utility at point of care [19-21]. To date, there has been limited detail on the performance characteristics of emerging RT-LAMP based tests in head-to-head comparisons with gold standard assays. Such information is critical to ensuring the safe deployment of new tests, given the clinical and public health consequences of erroneous test results.

Here we have shown the specific detection of SARS-CoV-2 RNA directly from clinical swabs without the need for RNA purification, using a LAMP assay targeting the N1 region of the SARS-CoV-2 N-gene in conjunction with the reverse transcriptase/GspSSD LF DNA polymerase mastermix from OptiGene Ltd. With a detection limit of 54 TCID_50_ mL^−1^ the N1-STOP-LAMP assay met each of the four FDA criteria for EUA. Based on reported data, as opposed to head-to-head comparisons, the N1-STOP-LAMP detection limit, is higher than several recently reported FDA-EUA COVID-19 tests. Expressed as virus copies per mL the N1-STOP-LAMP LoD is estimated at 54,000 virus copies per mL (assumes our TCID_50_ underestimates virus copy number by 1000-fold, see methods), compared to Cepheid GeneXpert Xpress 250 virus copies per mL or Abbott ID NOW COVID-19 test 125 virus copies per mL. Other EUA assays reporting LoDs similar to N1-STOP-LAMP include the Luminex ARIES SARS-CoV-2 assay (75,000 copies per mL) and the GenMark ePlex SARS-CoV-2 test (100,000 copies per mL). Given these performance metrics, and the test’s high positive predictive value (100%), we see N1-STOP-LAMP as well suited for widespread screening of large populations when COVID-19 prevalence is low. N1-STOP-LAMP could be used in large-scale, national testing programs to support aggressive COVID-19 contact tracing and disease suppression or elimination activities. The test could also play a role in near-point-of-care settings, such as outbreak investigation in hospitals and nursing homes, providing rapid-turnaround-time of results for health-care workers and highly vulnerable patients. The relatively simple assay format of N1-STOP-LAMP is also suitable for deployment in lower and middle-income countries, where access to sophisticated laboratory infrastructure is limited.

During this evaluation we noted opportunities for improvement of N1-STOP-LAMP. We observed that although 1 μL of UTM was tolerated in the OptiGene RT-LAMP formulation, larger quantities of sample matrix (UTM or PBS containing additional human material from the nasopharyngeal swab) were inhibitory to the RT-LAMP reaction. If more than 1 μL of spiked (purified virus or RNA as template) sample matrix was used in the assay, no amplification was observed (Fig. 1B). However, under ideal conditions (*i*.*e*., no sample inhibition from human components), the impact of using an increased template volume was shown, where 5 μL of UTM added directly to the N1-STOP-LAMP reaction was equal to the sensitivity of using purified RNA (Fig. 2B). Furthermore, addition of a simple RNA purification step, that does not require commercial kits, improved the detection sensitivity of N1-STOP-LAMP to rival RT-qPCR. Our follow-up of five false-negative results (Table 5, Fig. 3C), demonstrated the feasibility of including a simple, scalable and rapid magnetic-bead RNA purification step [26], with which we were able to detect SARS-CoV-2 RNA with N1-STOP-LAMP in 4/5 UTM specimens that had previously tested negative by the assay (Table S1). Other opportunities for improvement include the use of specific fluorescent probes to detect amplification products, thus facilitating multiplex reactions and the inclusion of an internal amplification control within each test [24,34].

As mentioned, recognised strengths of the LAMP assay include the rapid time to test result, with the majority of the positive clinical samples yielding a result in under 15 min, and the robust test format, testing directly from the swab eluate. Our preferred specimen type for N1-STOP-LAMP is a dry nasopharyngeal swab. Detailed examination of optimum swabs for the detection of influenza and other RNA viruses has shown dry swabs offer superior performance [35]. In the current study we have shown that N1-STOP-LAMP has satisfactory performance using dry swabs eluted in 1.5 mL of PBS (Fig. 3B); a format that potentially doubles the effective concentration of virus in the sample compared to 3 mL of UTM in the Copan format. Another strength of N1-STOP-LAMP is that the assay readily scales from small sample numbers (e.g. 8-well with portable detection unit) to high throughput (run with sample robotics and 96-well thermocyclers in a centralised laboratory).

While establishing the assay, we noted susceptibility to contamination that was easily addressed by the standard precautions used to prevent template carryover for any nucleic acid amplification test, such as: physical separation of mastermix preparation from sample inoculation; not opening post-amplification reaction tubes or plates; and inclusion of negative extraction controls. Addition of uracil-N-glycosylase has been reported as a strategy to limit the risk of amplicon contamination, however this is associated with a loss of detection sensitivity [36].

In this report we have shown N1-STOP-LAMP is robust diagnostic test for the specific and rapid detection of SARS-CoV-2. It is an alternative molecular test for SARS-CoV-2 that can be readily and seamlessly deployed, particularly when access to standard RT-qPCR based approaches are limited.

## Supporting information

Supplemental data file 1: GISAID acknowledgements

## 8. Author statements

### 8.1 Authors and contributors

Jean Lee:

Conceptualisation

Data curation

Formal analysis

Funding acquisition

Investigation

Methodology

Project administration

Resources

Software

Supervision

Validation

Visualisation

Writing - original draft

Writing - review & editing

Nickala Best:

Conceptualisation

Data curation

Formal analysis

Investigation

Methodology

Resources

Validation

Visualisation

Writing - original draft

Writing - review & editing

Julie McAuley:

Conceptualisation

Data curation

Formal analysis

Investigation

Methodology

Resources

Validation

Writing - review & editing

Jessica Porter:

Data curation

Formal analysis

Investigation

Methodology

Writing - review & editing

Torsten Seemann:

Conceptualisation

Investigation

Methodology

Software

Mark Schultz:

Conceptualisation

Investigation

Methodology

Software

Michelle Sait:

Formal analysis

Investigation

Methodology

Software

Nicole Orlando:

Formal analysis

Investigation

Methodology

Karolina Mercoulia:

Data curation

Investigation

Methodology

Susan Ballard:

Conceptualisation

Data curation

Formal analysis

Investigation

Methodology

Julian Druce:

Conceptualisation

Data curation

Formal analysis

Investigation

Methodology

Thomas Tran:

Investigation

Methodology

Mike Catton:

Conceptualisation

Investigation

Supervision

Writing - review & editing

Melinda Pryor:

Conceptualisation

Investigation

Methodology

Supervision

Writing - review & editing

Huanhuan Cui:

Investigation

Methodology

Writing - review & editing

Angela Luttick:

Investigation

Methodology

Writing - review & editing

Sean McDonald:

Conceptualisation

Formal analysis

Investigation

Methodology

Writing - review & editing

Arran Greenhalgh:

Conceptualisation

Formal analysis

Funding acquisition

Investigation

Methodology

Writing - review & editing

Jason Kwong:

Conceptualisation

Formal analysis

Funding acquisition

Methodology

Writing - original draft

Writing - review & editing

Norelle Sherry:

Conceptualisation

Formal analysis

Investigation

Methodology

Validation

Writing - original draft

Writing - review & editing

Maryza Graham:

Conceptualisation

Formal analysis

Investigation

Methodology

Validation

Writing - review & editing

Tuyet Hoang:

Conceptualisation

Funding acquisition

Investigation

Methodology

Validation

Writing - review & editing

Marion Herisse:

Conceptualisation

Funding acquisition

Investigation

Methodology

Resources

Validation

Writing - review & editing

Sacha Pidot:

Investigation

Methodology

Resources

Writing - review & editing

Deborah Williamson:

Conceptualisation

Formal analysis

Funding acquisition

Investigation

Methodology

Writing - review & editing

Benjamin Howden:

Conceptualisation

Formal analysis

Funding acquisition

Investigation

Methodology

Project administration

Resources

Writing - original draft

Writing - review & editing

Ian Monk:

Conceptualisation

Data curation

Formal analysis

Funding acquisition

Investigation

Methodology

Project administration

Resources

Validation

Writing - original draft

Writing - review & editing

Timothy Stinear:

Conceptualisation

Data curation

Formal analysis

Funding acquisition

Investigation

Methodology

Project administration

Resources

Software

Supervision

Validation

Visualisation

Writing - original draft

Writing - review & editing

## 8.4 Ethical approval

This study was conducted in accordance with the *National Health and Medical Research Council of Australia National Statement for Ethical Conduct in Human Research* 2007 (Updated 2018). The study was exempt from requiring specific approvals, as it involved the use of existing collections of data or records that contained non-identifiable data about human beings [37].

## Supplementary data

**Table S1.**
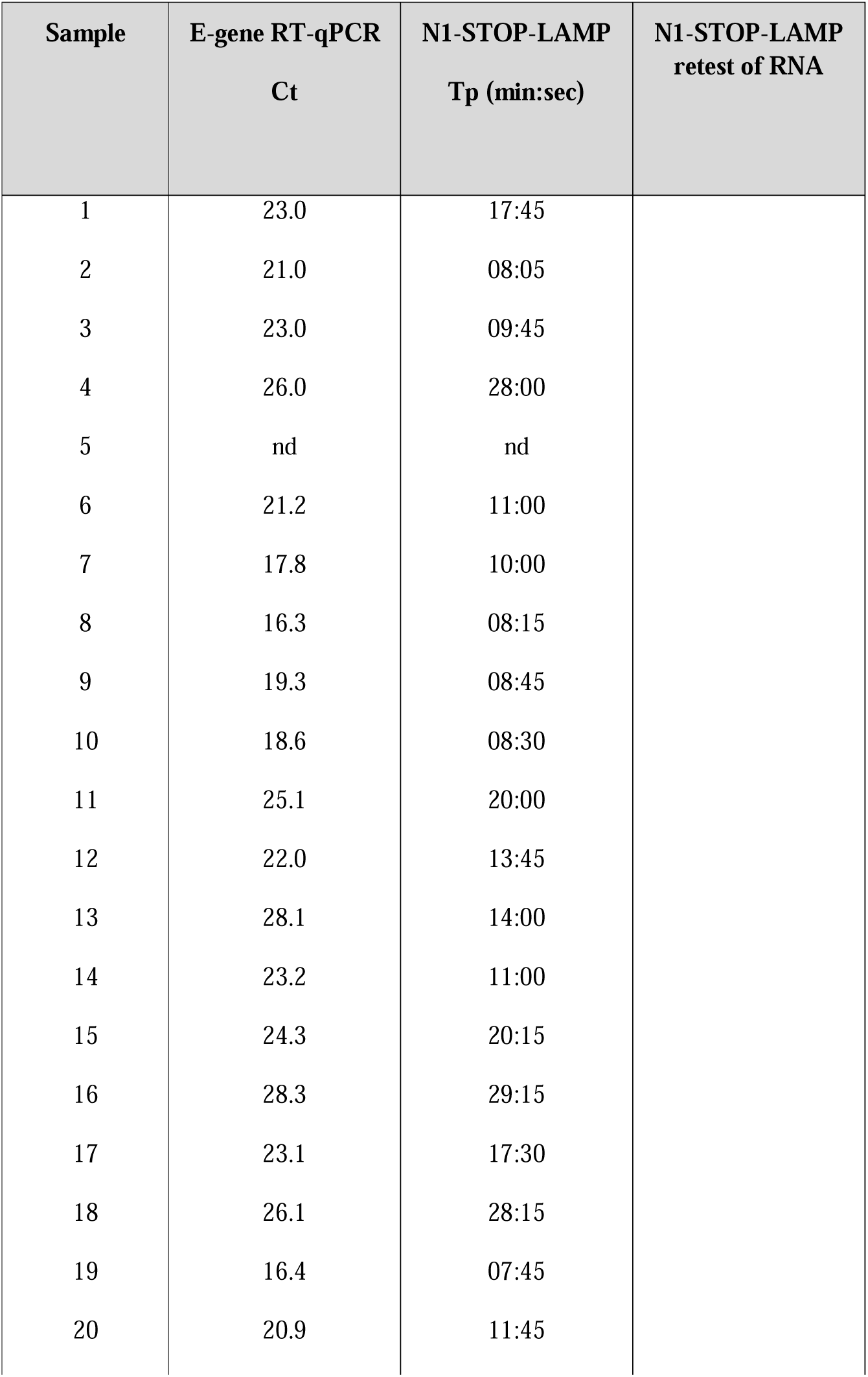
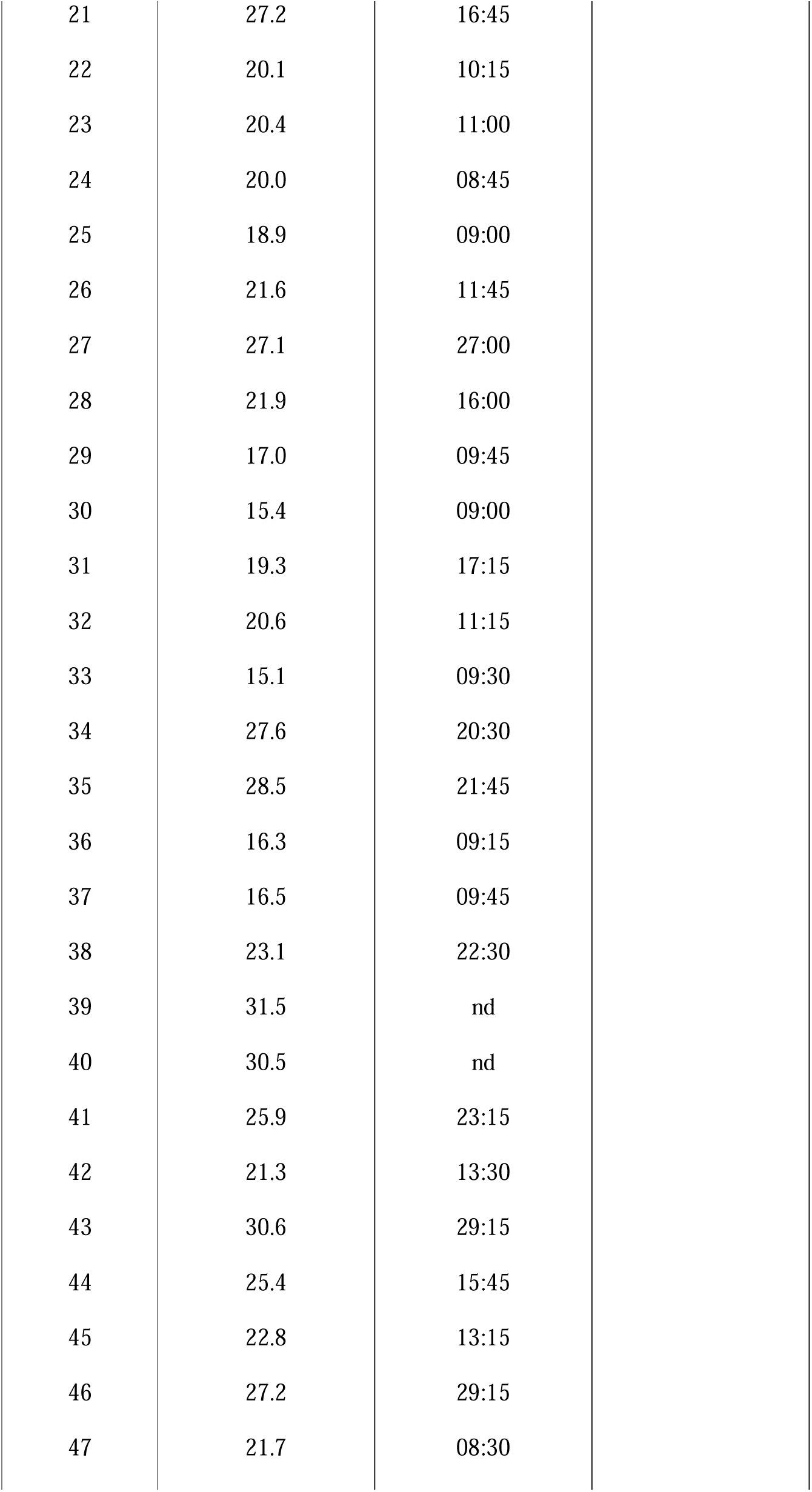
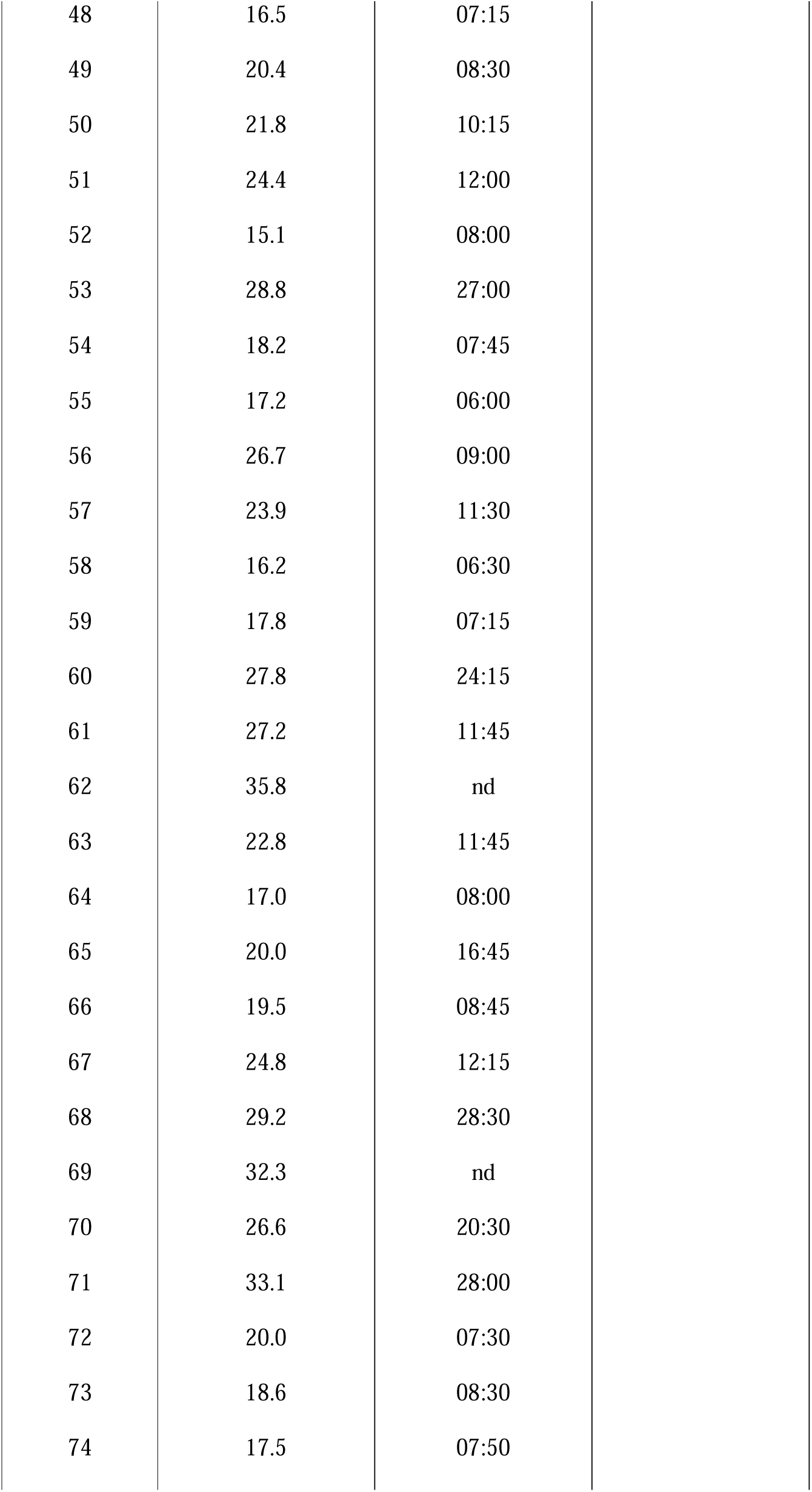
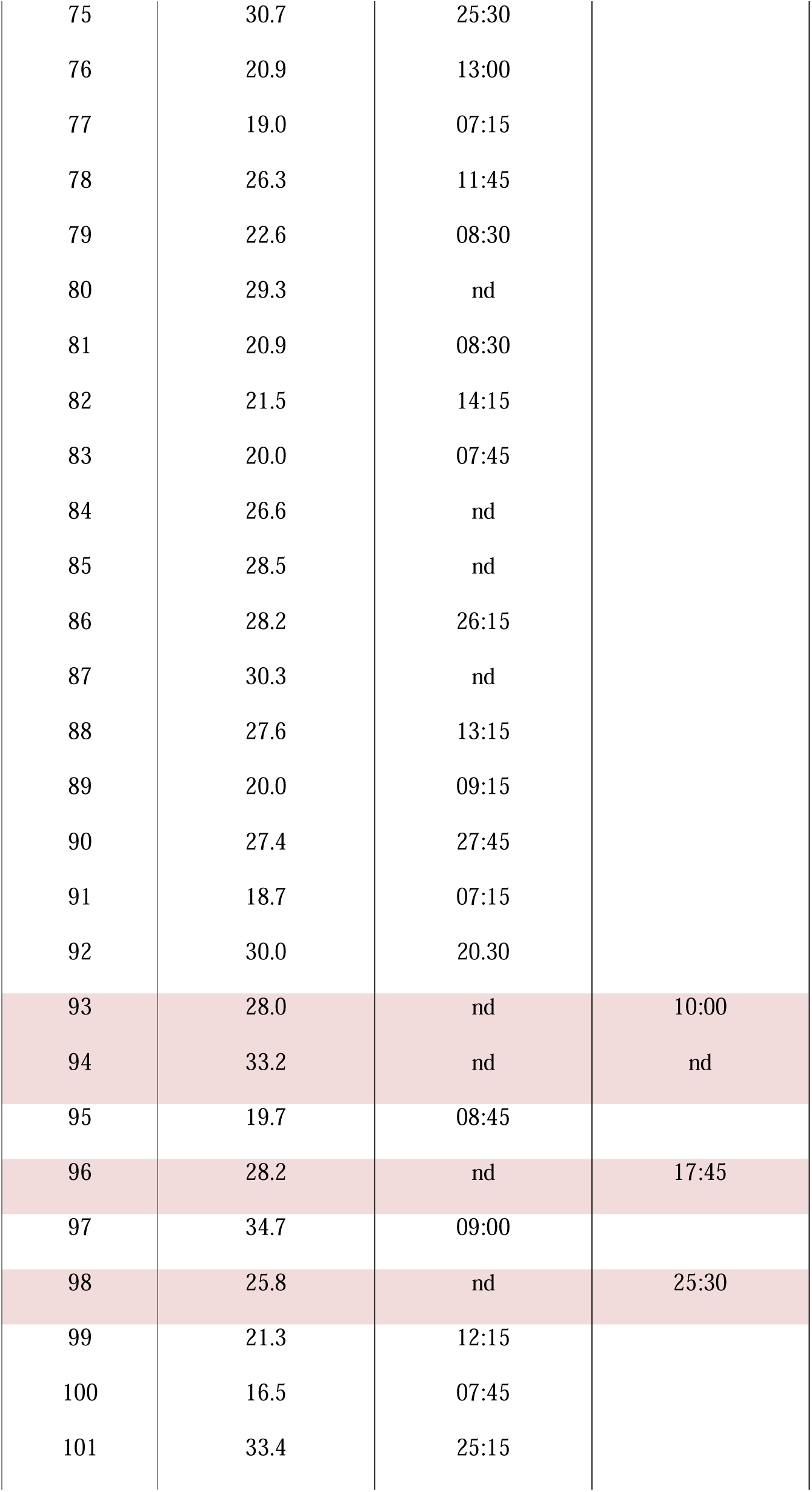
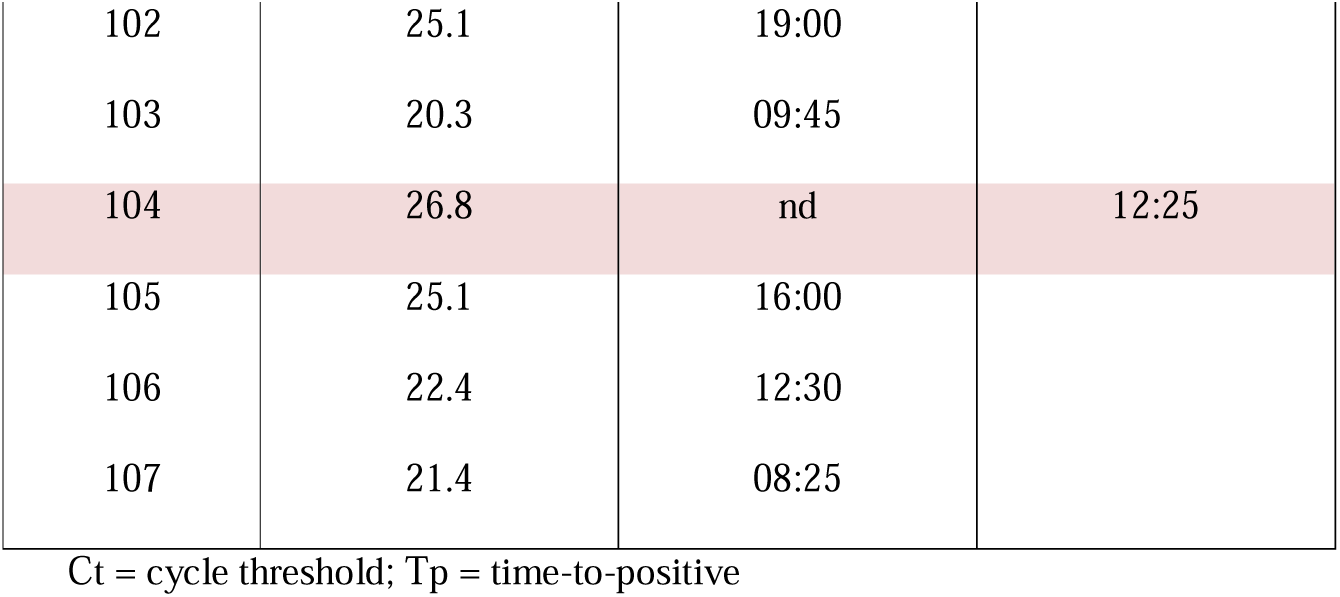
Clinical data positivity summary (107 E-gene RT-qPCR positives vs N1-STOP-LAMP Tp).

| Sample | E-gene RT-qPCR<br>Ct | N1-STOP-LAMP<br>Tp (min:sec) | N1-STOP-LAMP<br>retest of RNA |
| --- | --- | --- | --- |
| 1 | 23.0 | 17:45 |  |
| 2 | 21.0 | 08:05 |  |
| 3 | 23.0 | 09:45 |  |
| 4 | 26.0 | 28:00 |  |
| 5 | nd | nd |  |
| 6 | 21.2 | 11:00 |  |
| 7 | 17.8 | 10:00 |  |
| 8 | 16.3 | 08:15 |  |
| 9 | 19.3 | 08:45 |  |
| 10 | 18.6 | 08:30 |  |
| 11 | 25.1 | 20:00 |  |
| 12 | 22.0 | 13:45 |  |
| 13 | 28.1 | 14:00 |  |
| 14 | 23.2 | 11:00 |  |
| 15 | 24.3 | 20:15 |  |
| 16 | 28.3 | 29:15 |  |
| 17 | 23.1 | 17:30 |  |
| 18 | 26.1 | 28:15 |  |
| 19 | 16.4 | 07:45 |  |
| 20 | 20.9 | 11:45 |  |

|  |  |  |
| --- | --- | --- |
| 21 | 27.2 | 16:45 |
| 22 | 20.1 | 10:15 |
| 23 | 20.4 | 11:00 |
| 24 | 20.0 | 08:45 |
| 25 | 18.9 | 09:00 |
| 26 | 21.6 | 11:45 |
| 27 | 27.1 | 27:00 |
| 28 | 21.9 | 16:00 |
| 29 | 17.0 | 09:45 |
| 30 | 15.4 | 09:00 |
| 31 | 19.3 | 17:15 |
| 32 | 20.6 | 11:15 |
| 33 | 15.1 | 09:30 |
| 34 | 27.6 | 20:30 |
| 35 | 28.5 | 21:45 |
| 36 | 16.3 | 09:15 |
| 37 | 16.5 | 09:45 |
| 38 | 23.1 | 22:30 |
| 39 | 31.5 | nd |
| 40 | 30.5 | nd |
| 41 | 25.9 | 23:15 |
| 42 | 21.3 | 13:30 |
| 43 | 30.6 | 29:15 |
| 44 | 25.4 | 15:45 |
| 45 | 22.8 | 13:15 |
| 46 | 27.2 | 29:15 |
| 47 | 21.7 | 08:30 |
| 48 | 16.5 | 07:15 |
| 49 | 20.4 | 08:30 |
| 50 | 21.8 | 10:15 |
| 51 | 24.4 | 12:00 |
| 52 | 15.1 | 08:00 |
| 53 | 28.8 | 27:00 |
| 54 | 18.2 | 07:45 |
| 55 | 17.2 | 06:00 |
| 56 | 26.7 | 09:00 |
| 57 | 23.9 | 11:30 |
| 58 | 16.2 | 06:30 |
| 59 | 17.8 | 07:15 |
| 60 | 27.8 | 24:15 |
| 61 | 27.2 | 11:45 |
| 62 | 35.8 | nd |
| 63 | 22.8 | 11:45 |
| 64 | 17.0 | 08:00 |
| 65 | 20.0 | 16:45 |
| 66 | 19.5 | 08:45 |
| 67 | 24.8 | 12:15 |
| 68 | 29.2 | 28:30 |
| 69 | 32.3 | nd |
| 70 | 26.6 | 20:30 |
| 71 | 33.1 | 28:00 |
| 72 | 20.0 | 07:30 |
| 73 | 18.6 | 08:30 |
| 74 | 17.5 | 07:50 |
| 75 | 30.7 | 25:30 |  |
| 76 | 20.9 | 13:00 |  |
| 77 | 19.0 | 07:15 |  |
| 78 | 26.3 | 11:45 |  |
| 79 | 22.6 | 08:30 |  |
| 80 | 29.3 | nd |  |
| 81 | 20.9 | 08:30 |  |
| 82 | 21.5 | 14:15 |  |
| 83 | 20.0 | 07:45 |  |
| 84 | 26.6 | nd |  |
| 85 | 28.5 | nd |  |
| 86 | 28.2 | 26:15 |  |
| 87 | 30.3 | nd |  |
| 88 | 27.6 | 13:15 |  |
| 89 | 20.0 | 09:15 |  |
| 90 | 27.4 | 27:45 |  |
| 91 | 18.7 | 07:15 |  |
| 92 | 30.0 | 20:30 |  |
| 93 | 28.0 | nd | 10:00 |
| 94 | 33.2 | nd | nd |
| 95 | 19.7 | 08:45 |  |
| 96 | 28.2 | nd | 17:45 |
| 97 | 34.7 | 09:00 |  |
| 98 | 25.8 | nd | 25:30 |
| 99 | 21.3 | 12:15 |  |
| 100 | 16.5 | 07:45 |  |
| 101 | 33.4 | 25:15 |  |
| 102 | 25.1 | 19:00 |  |
| 103 | 20.3 | 09:45 |  |
| 104 | 26.8 | nd | 12:25 |
| 105 | 25.1 | 16:00 |  |
| 106 | 22.4 | 12:30 |  |
| 107 | 21.4 | 08:25 |  |

Ct = cycle threshold; Tp = time-to-positive

**Supplementary Table S2.**
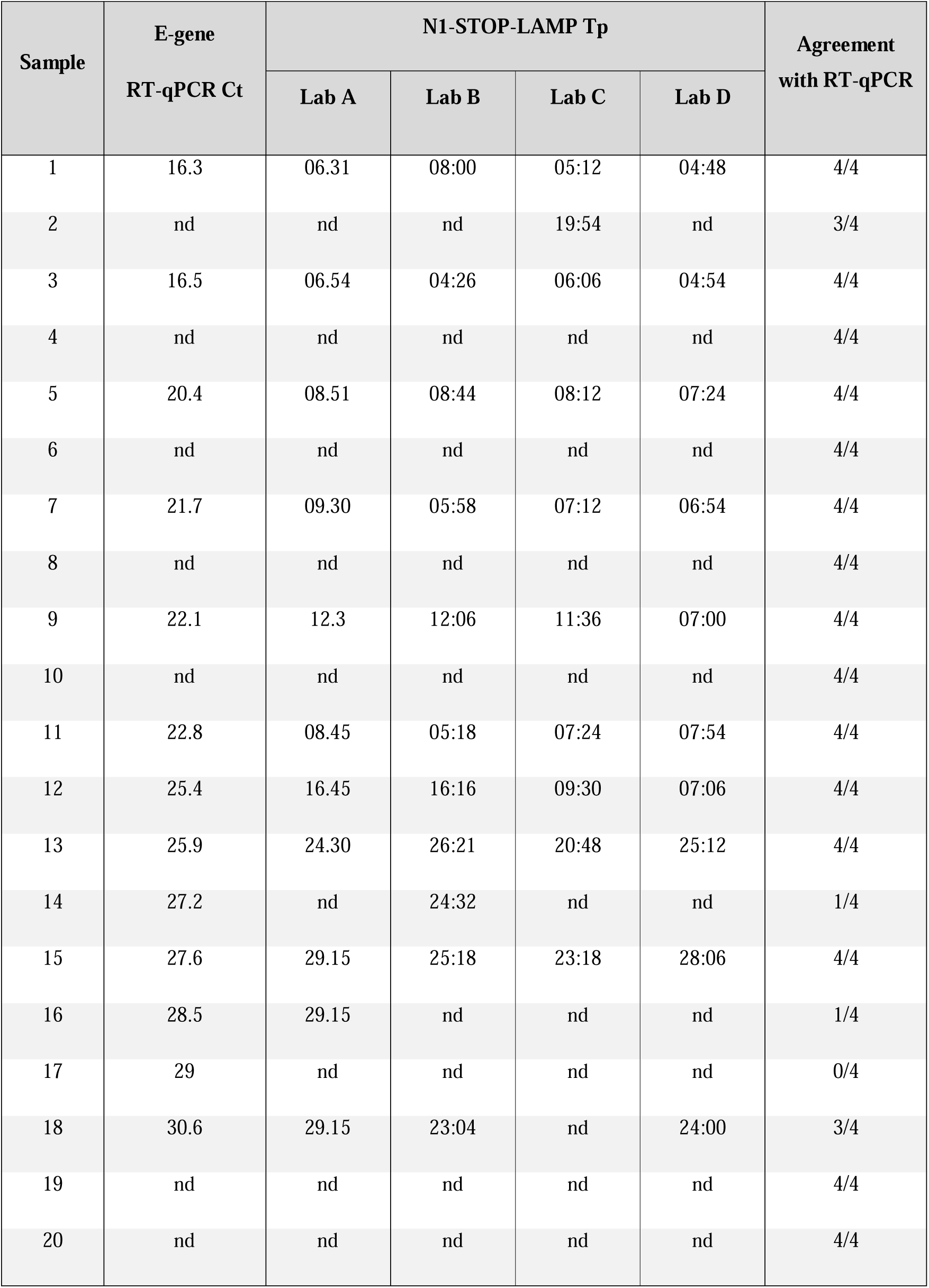
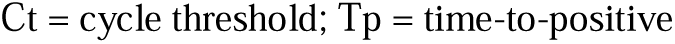
Results of N1-STOP-LAMP external quality assessment among four laboratories.

| Sample | E-gene | N1-STOP-LAMP Tp |  |  |  | Agreement<br>with RT-qPCR |
| --- | --- | --- | --- | --- | --- | --- |
|  | RT-qPCR Ct | Lab A | Lab B | Lab C | Lab D |  |
| 1 | 16.3 | 06.31 | 08:00 | 05:12 | 04:48 | 4/4 |
| 2 | nd | nd | nd | 19:54 | nd | 3/4 |
| 3 | 16.5 | 06.54 | 04:26 | 06:06 | 04:54 | 4/4 |
| 4 | nd | nd | nd | nd | nd | 4/4 |
| 5 | 20.4 | 08.51 | 08:44 | 08:12 | 07:24 | 4/4 |
| 6 | nd | nd | nd | nd | nd | 4/4 |
| 7 | 21.7 | 09.30 | 05:58 | 07:12 | 06:54 | 4/4 |
| 8 | nd | nd | nd | nd | nd | 4/4 |
| 9 | 22.1 | 12.3 | 12:06 | 11:36 | 07:00 | 4/4 |
| 10 | nd | nd | nd | nd | nd | 4/4 |
| 11 | 22.8 | 08.45 | 05:18 | 07:24 | 07:54 | 4/4 |
| 12 | 25.4 | 16.45 | 16:16 | 09:30 | 07:06 | 4/4 |
| 13 | 25.9 | 24.30 | 26:21 | 20:48 | 25:12 | 4/4 |
| 14 | 27.2 | nd | 24:32 | nd | nd | 1/4 |
| 15 | 27.6 | 29.15 | 25:18 | 23:18 | 28:06 | 4/4 |
| 16 | 28.5 | 29.15 | nd | nd | nd | 1/4 |
| 17 | 29 | nd | nd | nd | nd | 0/4 |
| 18 | 30.6 | 29.15 | 23:04 | nd | 24:00 | 3/4 |
| 19 | nd | nd | nd | nd | nd | 4/4 |
| 20 | nd | nd | nd | nd | nd | 4/4 |

Ct = cycle threshold; Tp = time-to-positive

**Supplementary Table S3.** N1-STOP-LAMP screen of NATtrol™ Respiratory Panel 2 (RP2) Controls.

| Virus/Organism | Strain | RP2 | RP2 | N1-STOP-LAMP |
| --- | --- | --- | --- | --- |
|  |  | Control 1 | Control 2 |  |
| Adenovirus Type 1 | N/A | Positive | Negative | Negative |
| Adenovirus Type 3 | N/A | Positive | Negative | Negative |
| Adenovirus Type 31 | N/A | Positive | Negative | Negative |
| *C. pneumoniae* | CWL-029 | Positive | Negative | Negative |
| Influenza A 2009 H1N1pdm | A/NY/02/2009 | Positive | Negative | Negative |
| Influenza A 2009 H3N2 | A/Brisbane/10/07 | Positive | Negative | Negative |
| Human Metapneumovirus Type 8 | Peru6-2003 | Positive | Negative | Negative |
| *M. pneumoniae* | M129 | Positive | Negative | Negative |
| Parainfluenza Type 1 | N/A | Positive | Negative | Negative |
| Parainfluenza Type 4 | N/A | Positive | Negative | Negative |
| Rhinovirus Type 1A | N/A | Positive | Negative | Negative |
| *B. parapertussis* | A747 | Negative | Positive | Negative |
| *B. pertussis* | A639 | Negative | Positive | Negative |
| Coronavirus 229E | N/A | Negative | Positive | Negative |
| Coronavirus HKU-1 | Recombinant | Negative | Positive | Negative |
| Coronavirus NL63 | N/A | Negative | Positive | Negative |
| Coronavirus OC43 | N/A | Negative | Positive | Negative |
| Influenza A H1N1 | A/New Cal/20/99 | Negative | Positive | Negative |
| Influenza B | B/Florida/02/06 | Negative | Positive | Negative |
| **Parainfluenza Type 2** | N/A | Negative | Positive | Negative |
| **Parainfluenza Type 3** | N/A | Negative | Positive | Negative |
| **RSV Type A** | 2006 isolate | Negative | Positive | Negative |

**Supplementary data file 1**. Global initiative on sharing all influenza data (GISAID) submitters table.

